# Profiling of urine bacterial DNA to identify an “oncobiome” in a mouse model of bladder cancer

**DOI:** 10.1101/364000

**Authors:** Catherine S. Forster, James J. Cody, Nirad Banskota, Crystal Stroud, Yi-Ju Hsieh, Dannah Farah, Olivia Lamanna, Olfat Hammam, Ljubica Caldovic, Michael H. Hsieh

## Abstract

**Introduction:** Recurrent urinary tract infections have been linked to increased risk of bladder cancer, suggesting a potential role of the urinary microbiome in bladder cancer pathogenesis.

**Objective:** Compare the urinary microbiomes in mice with and without bladder.

**Methods:** Longitudinal study of mice exposed to a dilute bladder-specific carcinogen (0.05% n-butyl-n-(4-hydroxybutyl) nitrosamine, BBN mice, n=10), and control mice (n=10). Urine was sampled monthly from individual mice for 4 months. Microbial DNA was extracted from the urine, and the V4 region of the 16S rRNA gene sequenced. Animals were sacrificed and their bladders harvested for histopathology. Bladder sections were graded by a blinded pathologist. The composition and diversity of the urinary microbiome were compared between the BBN and control mice. Metabolic pathway analysis was completed using PICRUST.

**Results:** Bladder histology in the BBN group showed normal tissue with inflammation (BBN-normal, n=5), precancerous pathologies, (BBN-precancerous, n=3), and invasive cancer (BBN-cancer, n=2). Alpha diversity did not differ between the mice exposed to BBN and the control mice at any timepoint. There were no differences in the urinary microbiomes between the BBN and control mice at baseline. At month 4, mice exposed to BBN had higher proportion of both *Gardnerella* and *Bifidobacterium* compared to control mice. There were no differences in proportions of specific bacteria between either the BBN-precancer or BBN-cancer and controls at month 4. However, the BBN-normal mice had higher proportions of *Gardnerella*, *Haemophilus*, *Bifidobacterium*, and *Ureaplasma Actinobaculum*, and lower proportions of *Actinomyces*, compared to control mice at month 4. Functional pathway analysis demonstrated increases in genes related to purine metabolism, phosphotransferase systems, peptidases, protein folding, and bacterial toxins in the BBN-mice compared to control mice at month 4.

**Conclusion:** Mice exposed to 4 months of BBN, a bladder-specific carcinogen, have distinct urine microbial profiles compared to control mice.

## Introduction

Bladder cancer is the 4^th^ most common malignancy among men in the U.S. [1] However, despite the high prevalence of bladder cancer, death rates from this neoplasm have decreased only slightly for women and not at all for men. These mortality rates are in part driven by late initial diagnosis of advanced bladder cancer, and poor ability to predict which patients with originally non-muscle-invasive cancer will progress to muscle-invasive disease. The ability to identify patients who are at risk for cancer, or progression of disease, through non-invasive means would have a significant impact in this population.

A number of risk factors for the development of bladder cancer have been identified, including an association between urinary tract infections (UTI) and bladder cancer. While there is a well-established causative relationship between infections with the urogenital parasite *Schistosoma haematobium* and bladder cancer,[5] [6] the association between bacterial urinary tract infections and bladder cancer is less clear. Data from large epidemiological studies suggest that an association may be present,[7][8] with other work suggesting that recurrent UTIs may be a particular risk factor for squamous cell carcinoma of the bladder [11]. However, other work has not found such an relationship between UTI and bladder cancer.[9][10] Conversely, the presence of asymptomatic bacteriuria has been associated with lower recurrence rate and longer disease-free survival in patients with non-muscle invasive bladder cancer.[12] The hypothesis behind this observed protective effect of asymptomatic bacteriuria is that activation of the immune system as a result of bacteriuria inhibits tumor formation.[13] Taken together, this suggests that immune responses to bacteria within the bladder may play a role in bladder oncogenesis.

Although urine has been classically considered to be sterile, technological advances have led to identification of a diverse community of bacteria within the bladder, known as the urinary microbiome [16–18]. Given the potential immune-associated effect of bacteriuria on bladder oncogenesis, we postulated that there may be specific patterns within the urinary microbiome that would exert either a protective or harmful effect on tumor development in those at risk for bladder cancer. Indeed, prior work focusing on the microbiomes of other organ systems have documented microbial changes in the setting of cancer, including oral squamous cell carcinoma,[19] cervical cancer,[20] and colon cancer.[21] Therefore, we sought to determine whether there were changes in the urinary microbiome in an experimental mouse model of bladder cancer, and whether we could identify distinct profiles associated with specific lesions along the spectrum of bladder oncogenesis. To test our hypothesis that mice with bladder cancer would harbor a distinct urinary “oncobiome”, we used an established model in which a dilute bladder-specific carcinogen (n-butyl-n-(4-hydroxybutyl) nitrosamine, BBN) is administered to mice in their drinking water. The BBN model, a widely used and well-established model of bladder carcinogenesis, [22,23] reliably leads to tumor formation in both mice and rats. Although the incidence of cancer is not 100%, this can be seen as an advantage given that an array of pathologies are obtained. Work by our group and others has shown that BBN-induced bladder cancers in mice closely resemble human cancers, both histologically [24], and by gene expression analysis, with a particular resemblance to muscle-invasive disease noted by several groups [25–27].

## Methods

### Mice

Twenty female C57BL/6 mice (received at 5 weeks of age from Jackson Laboratories, Bar Harbor, ME) were used in this work. Ten mice were treated with the bladder-specific carcinogen BBN (n-butyl-n-(4-hydroxybutyl) nitrosamine, Sigma-Aldrich, St. Louis, MO) at 0.05% in their drinking water *ad libitum* (tap water filtered through a Milli-Q Academic system, MilliporeSigma, Burlington, MA) over a period of five months, and ten mice were given unmanipulated tap water (municipal water supply, Rockville, MD). All ten BBN-treated mice were housed together in a single cage, as were the ten control mice. The bedding (Aspen Chip^®^, Northeast Products Corporation, Warrensburg, NY) and chow (LabDiet^®^ Prolab^®^ RMH 1000, PMI Nutrition, Brentwood, MO) were the same between the two groups. Mice were housed in adjacent cages in the same holding room of the animal facility.

### Urine Collection

Urine samples were collected at baseline, beginning 1 week after arrival to allow for acclimation to our facility, and then at monthly intervals following the initiation of exposure to BBN. At each time point, urine samples were collected from individual mice over the course of three days. At each time point, urine samples ranging from 10 – 100 μl were collected from each mouse. Parafilm^®^ (Bemis Company, Inc., Oshkosh, WI) was used to cover a tube rack such that the sterile surface of the parafilm was exposed. Mice were then placed on the sterile surface and pressure was placed on the lower abdominal area until urination occurred. Mice were limited from walking around to avoid contamination of the parafilm, and any samples contaminated with feces were discarded. Urine was collected from the clean areas of the parafilm using a sterile barrier tip. Following collection, samples were stored at −80 °C until the completion of the study, at which point all the samples were processed as a single batch.

### Microbiome Analysis

DNA was isolated from mouse urine using Qiagen DNeasy Powersoil Kit (Hilden, Germany); bacterial DNA in each sample was quantified using Femto Bacterial DNA Quantification kits (Zymo Research, Irvine, CA) to determine the fraction of bacterial DNA in each DNA sample. V4 regions of 16S rRNA genes were amplified using primers 5’-TCGTCGGCAGCGTCAGATGTGTATAAGAGACAGGTGCCAGCMGCCGCGGTAA-3’ and 5’-GTCTCGTGGGCTCGGAGATGTGTATAAGAGACAGGACTACHVGGGTWTCTAAT-3’ (IDT DNA, Coralville, IA) and the following reagent concentrations: 600 mM Tris-SO_4_ (pH 8.9), 180 mM (NH_4_)_2_ SO_4_, 20 mM MgSO_4_, 2mM dGTP,2mM dTTP, 2nM dCTP, 10% glycerol, and thermostable AccuPrime protein (Thermo Fisher Scientific, Waltham, MA) and 25 ng of template DNA in 20 μl total volume. Amplification conditions were 2 minutes at 95°C initial denaturation followed by 28 cycles of 20 seconds denaturation at 95°C, 15 seconds annealing at 55°C and a 5-minute extension at 72°C, and a 5-minute final extension at 72°C. Amplification products were purified with the AMPure XP system (Beckman Coulter Life Sciences Division, Indianapolis, IN) and quantified by a Qubit dsDNA assay (Thermo Fisher Scientific). Quality and size of amplification products were also verified with an Agilent 2100 Bioanalyzer kit (Agilent Technologies, Santa Clara, CA). Indexing and pooling of amplification products were carried out according to Illumina’s 16S Metagenomic Sequencing Library Preparation protocol. The resulting library was sequenced using Illumina MiSeq Reagent Kits v2 (500 cycles) at the Georgetown University Genomics and Epigenomics Shared Resource (Georgetown University, Washington, DC).

### Histology

Bladders were harvested from all BBN-exposed mice and two randomly chosen control mice after five months. Bladders were prepared for sectioning and staining by standard methods, briefly summarized here. Tissues were fixed in 10% formalin, dehydrated through a graded series of alcohols, embedded in paraffin, and stained with haematoxylin and eosin. Tissue sectioning and staining was performed by the Research Pathology Core Laboratory at the George Washington University (Washington, DC). Bladders were analyzed by a pathologist trained in bladder biology (OH), who assessed them in a blinded fashion.

### Statistical Analysis

After removing primer sequences present from FASTQ files using CutAdapt [28], these files were processed using Mothur [29]. In brief, the paired-end FASTQ reads were combined, after which ambiguous reads as well as reads longer than 275 bp were removed. We followed this step by merging all duplicate reads. These sequences were then aligned to the V4 region of the Silva reference database. Specifically, we used release 132 of the Silva reference database. The Silva file can be downloaded at this link: https://www.mothur.Org/w/images/3/32/Silva.nr_v132.tgz Chimeric sequences were then removed using the VSEARCH command [30]. Finally, our sequences were clustered using the opticlust algorithm [31]. The operational taxonomic units (OTUs) generated were then classified taxonomically. We followed the Miseq protocol available on https://www.mothur.org/wiki/MiSeqSOP.

For further analysis, we used STAMP [32]. The groups were primarily compared in a pairwise manner, using the Bonferroni correction for multiple comparisons. For statistical significance, we used Welch’s two-sided t-test and confidence intervals of 95%. Principal component analysis (PCA) plots along with extended error bar plots were generated using STAMP. We also used Phyloseq for further analysis[33]. Specifically, principal coordinate analysis (PCoA) plots were generated using Phyloseq and ggplot2[34]. For functional analysis, we used PICRUSt [35]. Data generated by PICRUSt was then loaded to STAMP for statistical analysis and chart generation.

## Results

In the BBN-treated group, a range of pathologies were observed. Of the ten mice who received BBN, five did not develop cancer, and had histology consistent with inflammation (“normal-like”),(Figure 1, panel b) three had either urothelial dysplasia, hyperplasia, or carcinoma in situ on histology (“precancer”),(Figure 1, panel c) and two developed invasive urothelial carcinomas (“cancer”), one of which had features of a squamous cell carcinoma.(Figure 1, panel d) The bladders of the control mice showed normal histology, as expected.(Figure 1, panel a)

**Figure 1.**
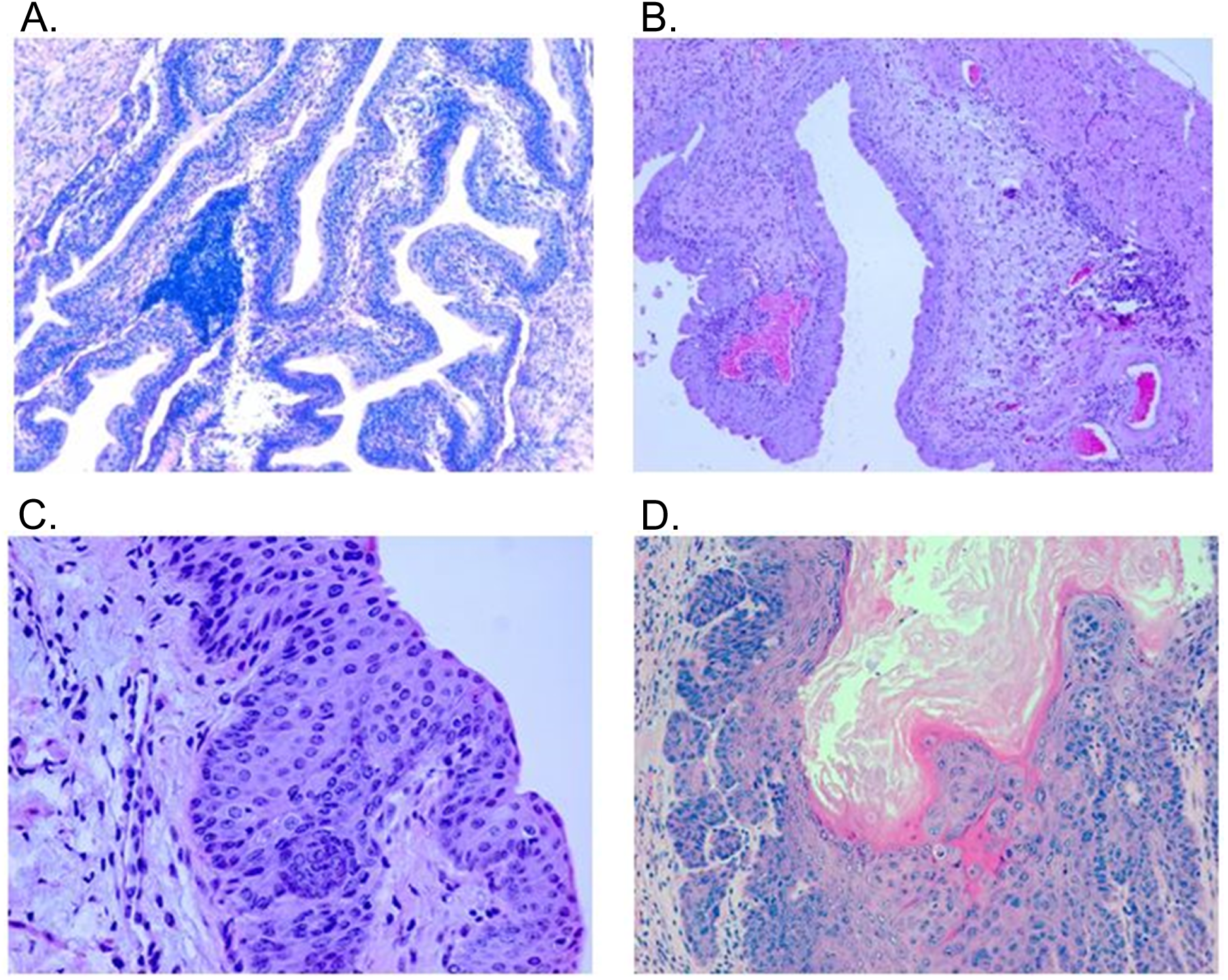
Bladder Histology of Control and BBN-Exposed Mice. Representative histology from BBN-treated and control mice. Bladders from control or BBN-treated mice were harvested after 5 months, and formalin-fixed paraffin-embedded tissue sections were stained with hematoxylin and eosin. Compared to the normal urothelium in the control mice (A, 100X), the BBN-treated mice exhibited a range of disease states from normallike with chronic inflammation (B, 100X) to dysplasia (C, 400X) to invasive bladder cancer with features of transitional cell and squamous cell carcinoma (D, 200X).

There was no significant difference in either the Shannon diversity index or the chao1 index between the aggregate BBN-treated mice and the control mice at either month 0 or month 4 (Figure 2A). There was also no difference in either the Shannon diversity index or the chao1 index between mice in either the BBN-normal, BBN-precancer, or BBN-cancer and the control mice at month 0 (Figure 2B) or month 4. (Figure 2C). Similarly, there was no change in either the Shannon diversity or Chao1 index for the between control mice at month 0 and month 4.

**Figure 2A:**
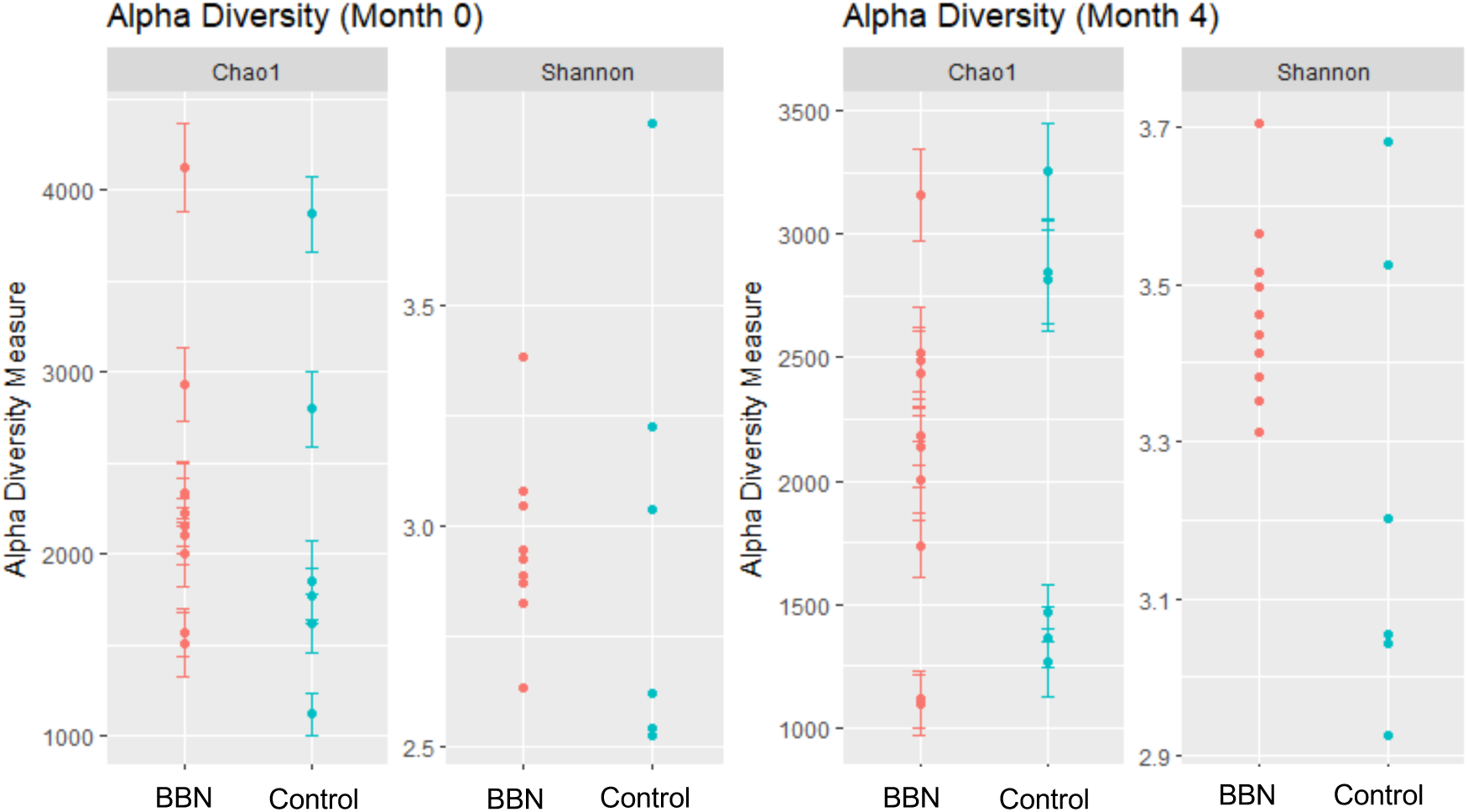
Alpha Diversity in BBN-treated and Control Mice. Comparison of indices of alpha diversity between groups. A) There is no difference in either the Shannon Diversity Index or Chaol Index between mice exposed to BBN (red) and control mice (blue) at month 0 or month 4. There is no difference between either the Shannon Diversity Index of the Chao1 Index between control mice (red) and the BBN-normal (green), BBN-pre-cancer (blue), or BBN-cancer (purple) at month 0 (panel B) or month 4 (panel C).

**Figure 2B.**
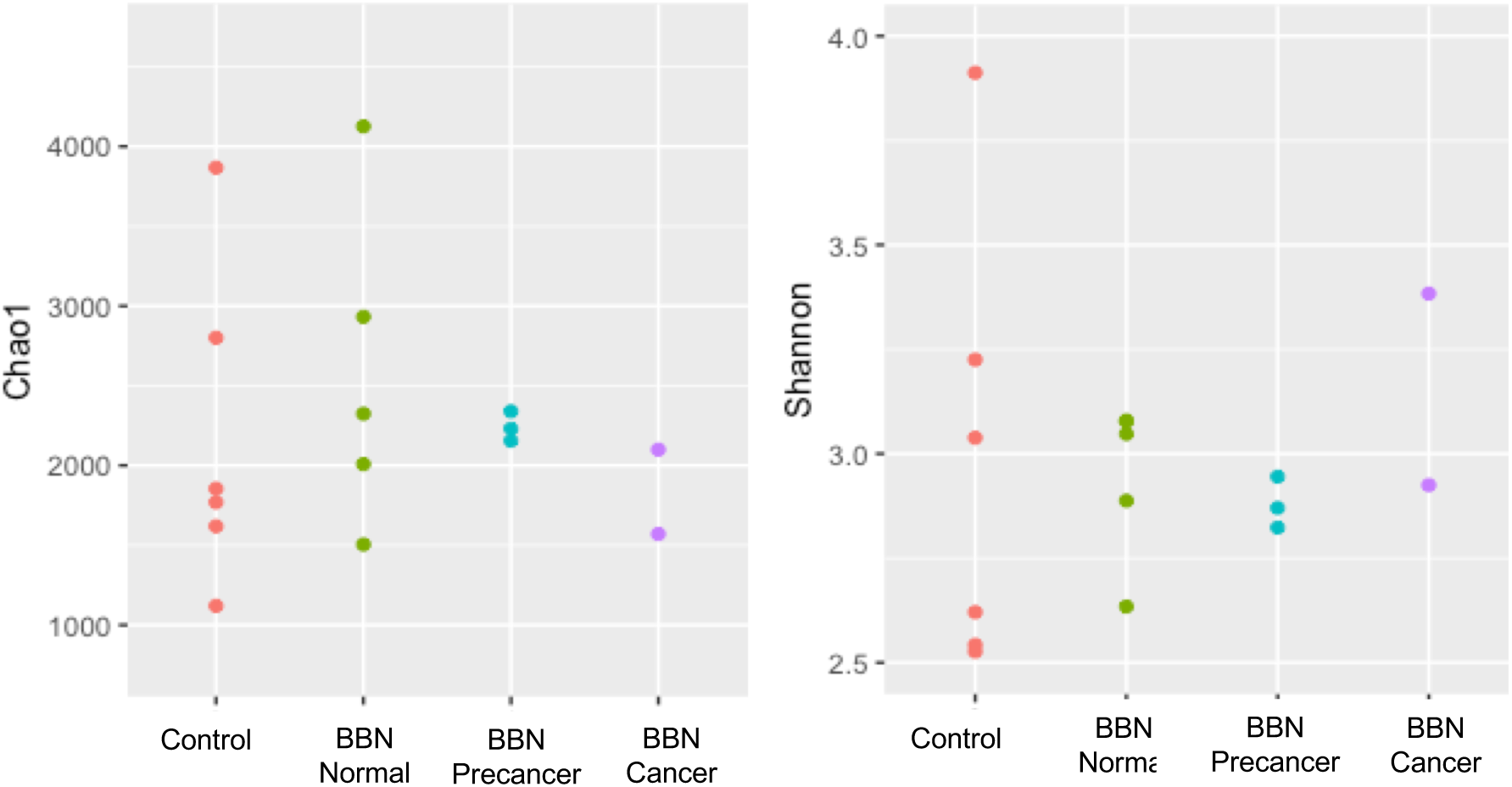
Alpha Diversity at Month 0 Between Sub-Groups.

**Figure 2C.**
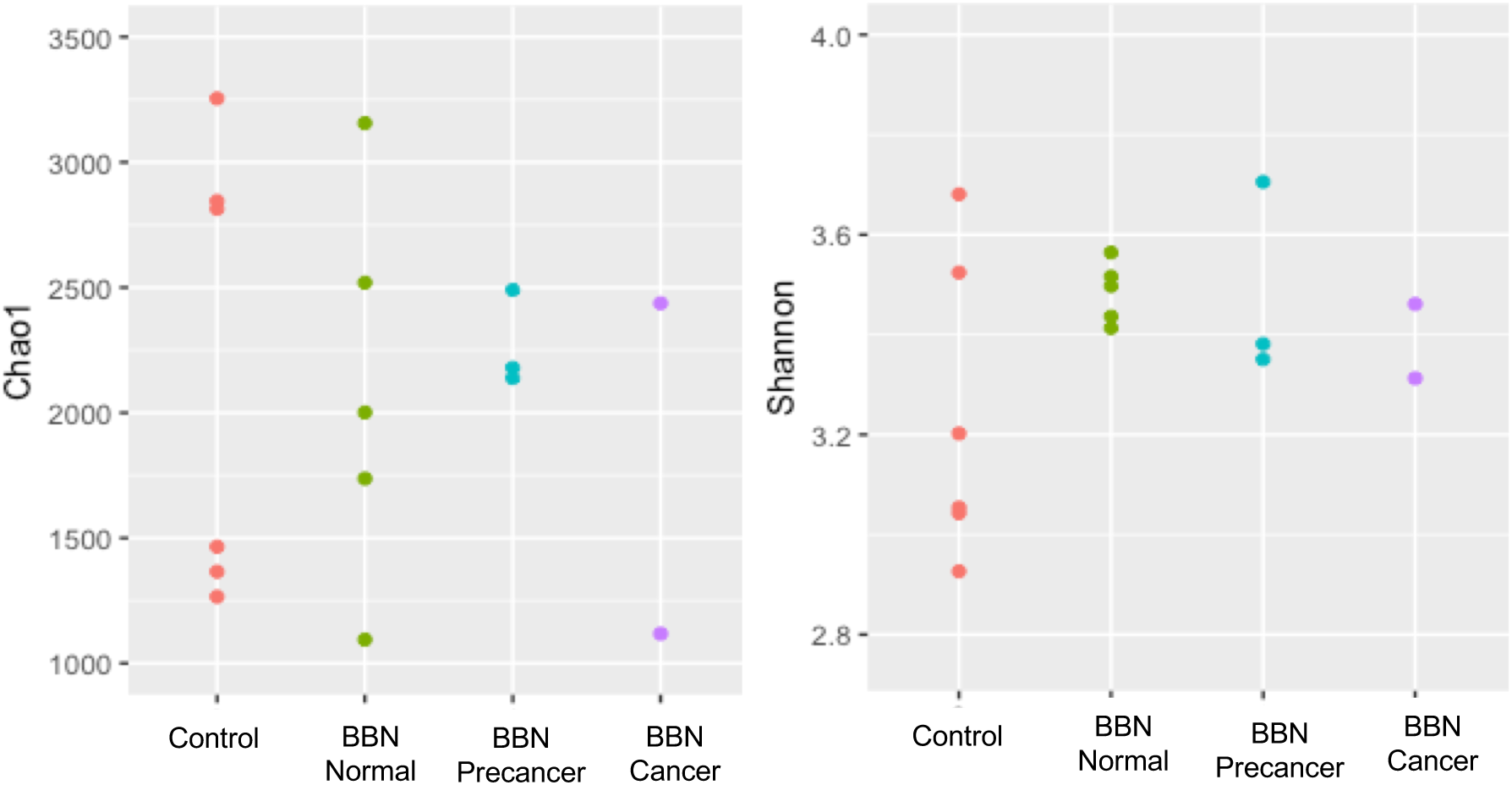
Alpha Diversity at Month 4 Between Sub-Groups.

Principal component analysis (PCA) demonstrated clustering of the microbiome for most mice at month 0, while there was divergence at month 4 between the BBN-treated mice and the controls (Figure 3). However, separate clusters correlating to histology was not observed. Of note, the one major outlier at month 0 went on to develop invasive carcinoma. The urinary microbiome of that mouse at month 0 was composed of different bacteria than the other mice in this work (Figure 4A). The most prevalent bacteria in the urinary microbiome of the outlying mouse were: *Rubellimicrobium*, *Escherichia*, *Roseococcus*, *Roseomonas*, *Kaistobacter*, and *Sphingomonas* (Figure 4B). Comparatively, the most common bacteria in the remaining urinary microbiomes were: *Escherichia*, *Prevotella*, *Veillonella*, *Streptococcus*, *Staphyloccoccus*, and *Neisseria.*

**Figure 3.**
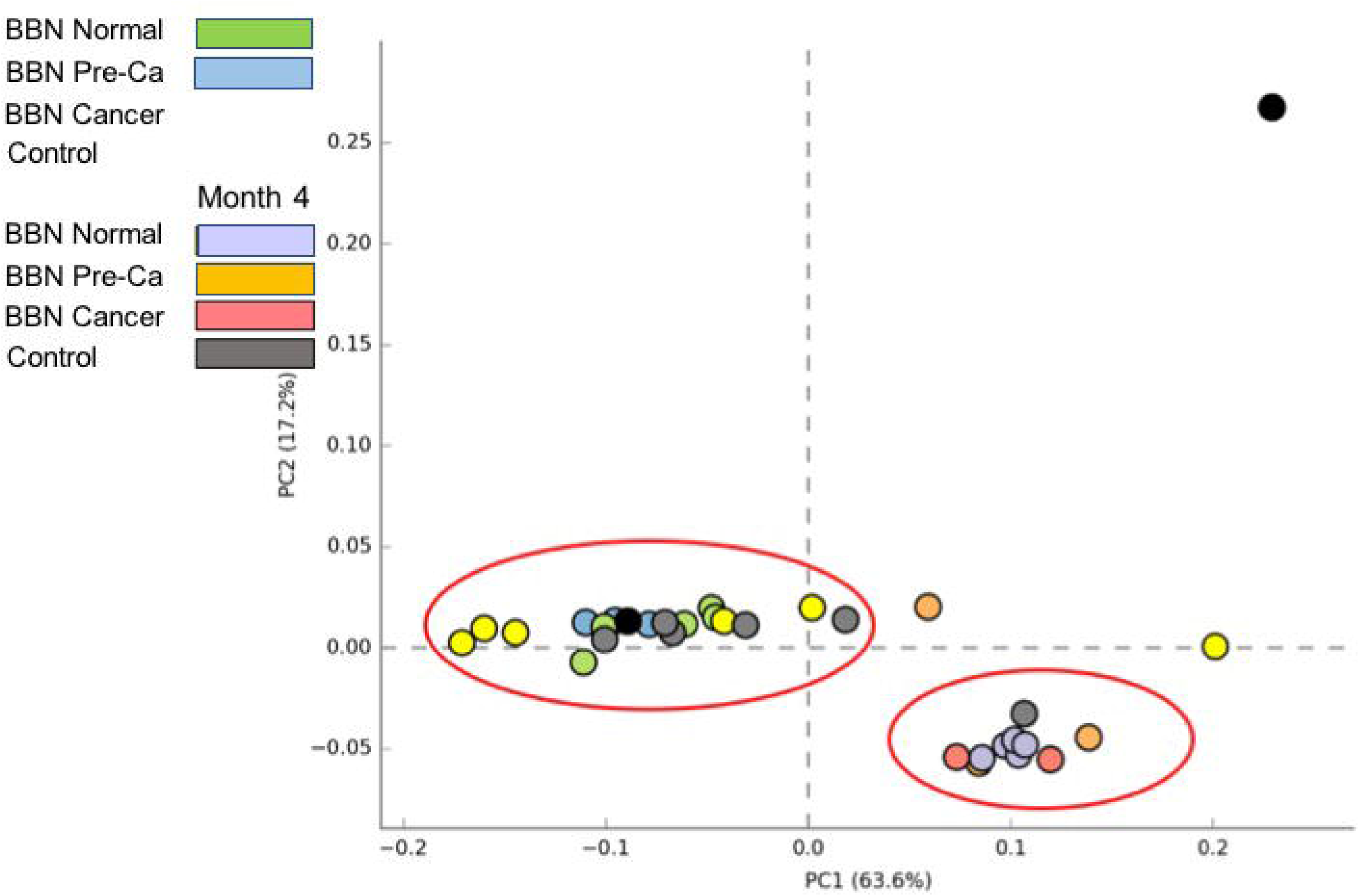
Principal component analysis (PCA) plots of urine “oncobiomes”. Urine samples were collected at baseline and at monthly intervals from control or BBN-treated mice over 4 months and the bacterial communities profiled by 16S v4 rRNA gene sequencing. At study completion, bladders were harvested and analyzed histologically, indicating a range of pathologies. BBN data sets were then subdivided by histological findings [BBN Normal (normal-like with inflammation, n = 5), BBN Pre-Ca (precancerous lesions, n = 3), BBN Cancer (invasive cancers, n = 2)] for comparison to control samples (n = 6). BBN and control samples largely clustered together at baseline (Month 0). However, by month 4, while the control samples still mostly clustered with the baseline samples, the BBN samples were predominantly grouped in a distinct cluster. An outlier at Month 0 (upper right of plot) later developed invasive carcinoma from BBN treatment. Plots were generated using STAMP (STatistical Analysis of Metagenomic Profiles).

**Figure 4.**
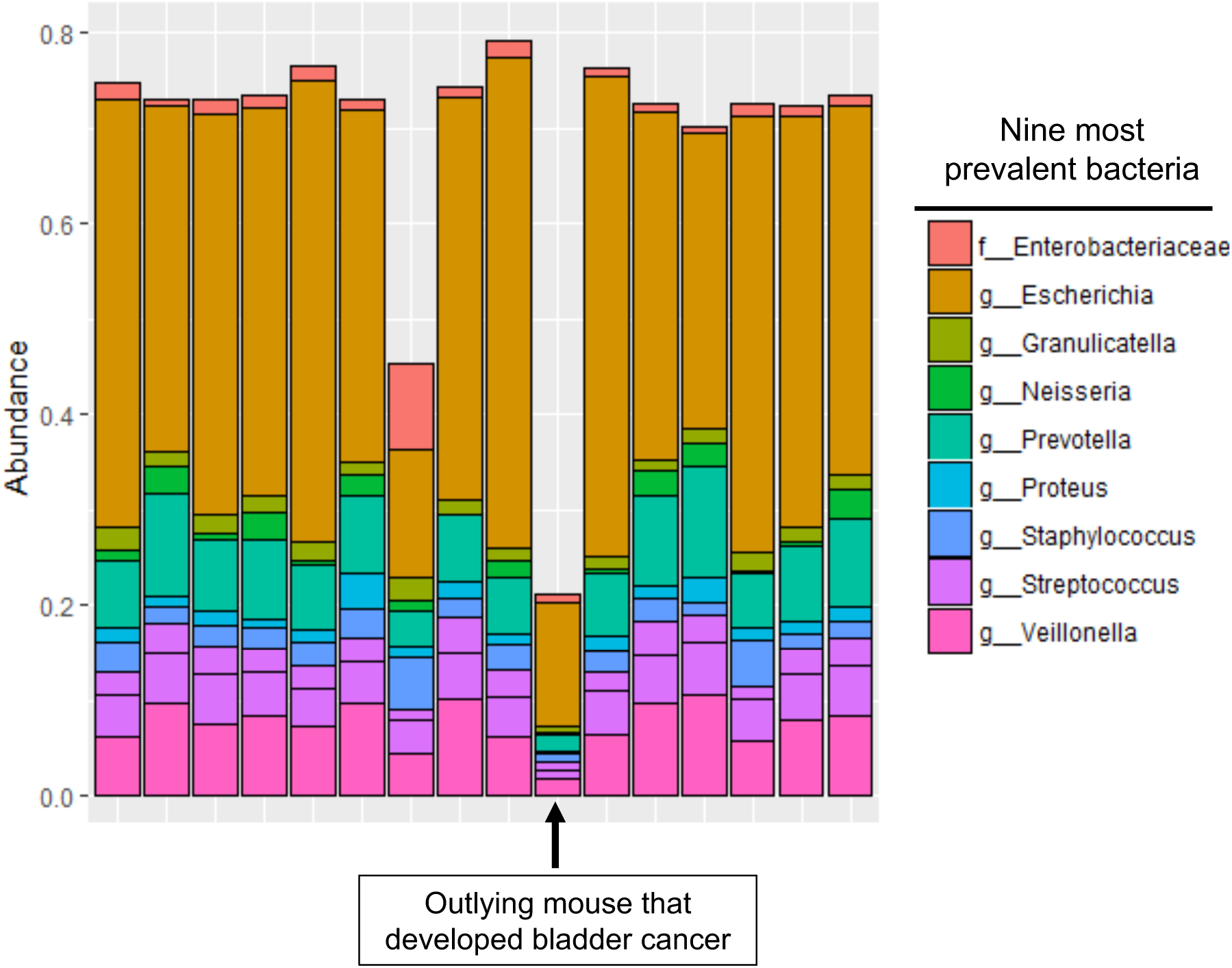
Relative Abundance of Bacteria in Urinary Microbiome at Month 0. Relative abundance charts. A) Relative abundancies of the nine most prevent bacteria in the urinary microbiome at month 0. The arrow indicates the mouse whose urinary microbiome at month 0 was an outlier on the PCA chart that developed cancer.

**Figure 4B:**
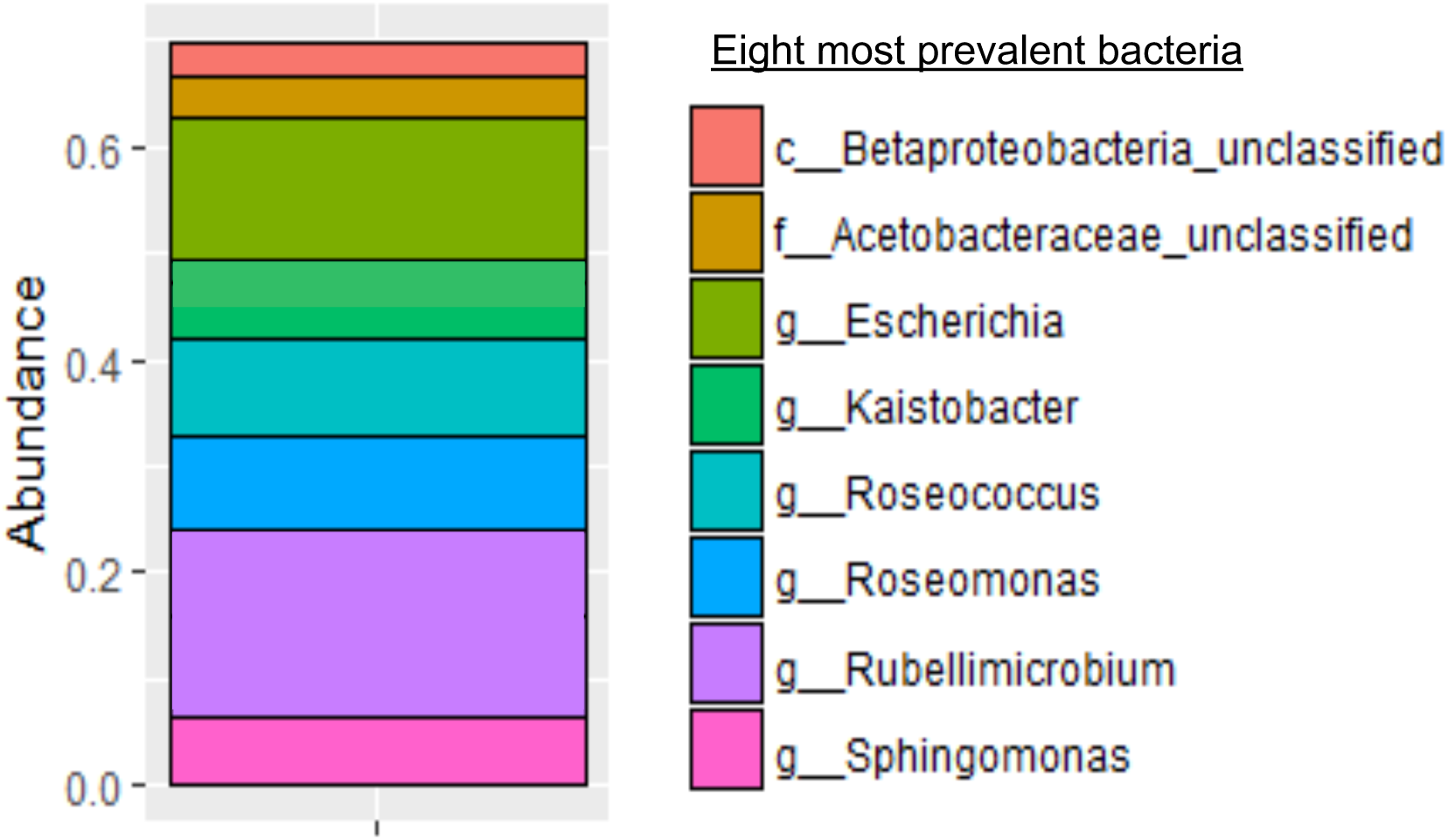
Relative Abundance of Bacteria in Urinary Microbiome at Month 0 of Outlying Mouse that Developed Bladder Cancer. B) Relative abundancy chart of the eight most common bacteria in the urinary microbiome of the outlying mouse at month 0.

There were no significant differences in proportions of specific bacteria at months 0 through month 3 between the aggregate BBN-treated (regardless of histology) and control mice. However, at month 4, the aggregate BBN-treated mice had significantly higher proportion of *Gardnerella* (corrected p-value = 0.047) and *Bifidobacterium* (corrected p-value 0.045) compared to the control mice. (Figure 5A) The corresponding functional analysis by PICRUST demonstrated that the urinary microbiome in the BBN-treated mice at month 4 had increases in multiple pathways compared to the control mice at month 4. A total of 44 pathways were differentially expressed, including: purine metabolism, phosphotransferase systems, peptidases, protein folding, and bacterial toxins. (Figure 6) There were no other differences noted in the functional analysis between any other months.

**Figure 5:**
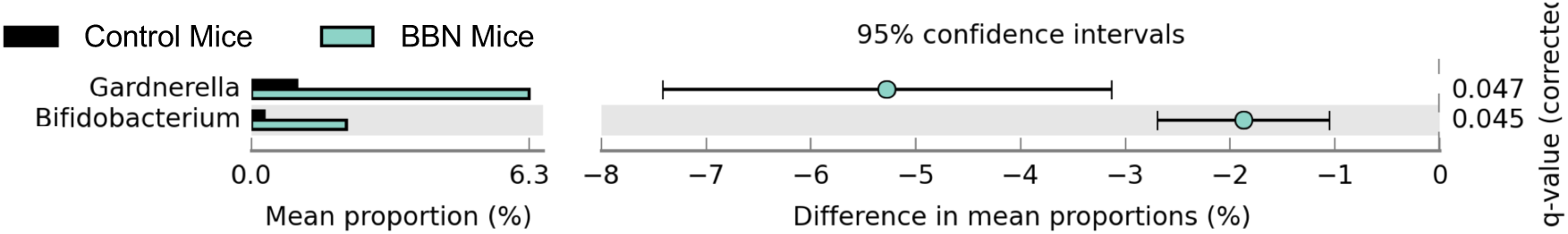
Mean Proportions Chart of BBN-Treated Versus Control Mice at Month 4. Mean proportion charts. A) Mean proportion chart comparing the proportion of the bacteria in the urinary microbiome of control (black) versus BBN-exposed (green) mice at month 4. Proportions of *Gardernella* and *Bifidobacterium* were significantly higher in the BBN-exposed month compared to the controls.

**Figure 5B:**
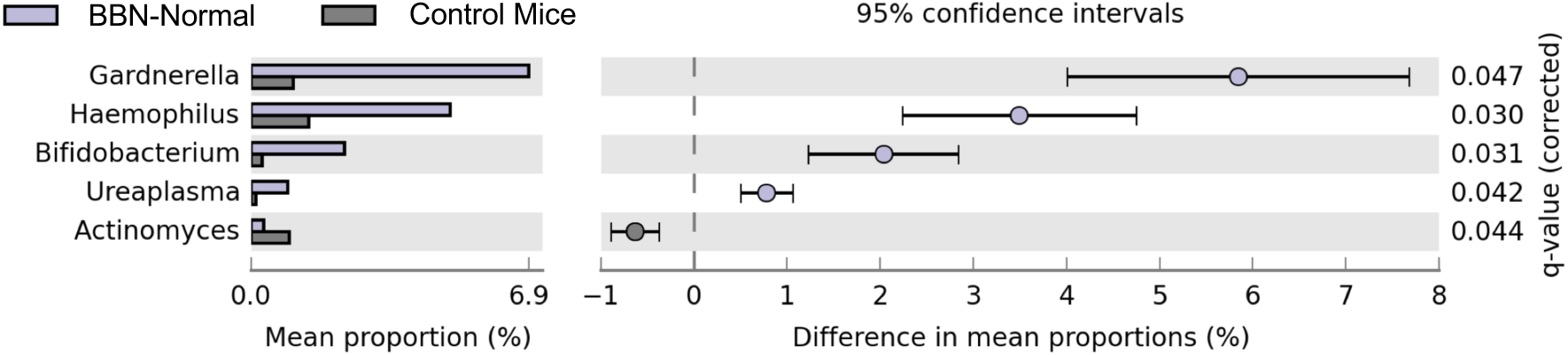
Mean Proportions Chart of BBN-Normal Versus Control Mice at Month 4. B) Mean proportion chart comparing the urinary microbiome of BBN-normal versus control mice at month 4, with higher proportions of *Gardernella*, *Haemophilus*, *Bifidobacterium*, and *Ureaplasma* in the BBN-normal mice, and *Actinomyces* in the control mice.

**Figure 6:**
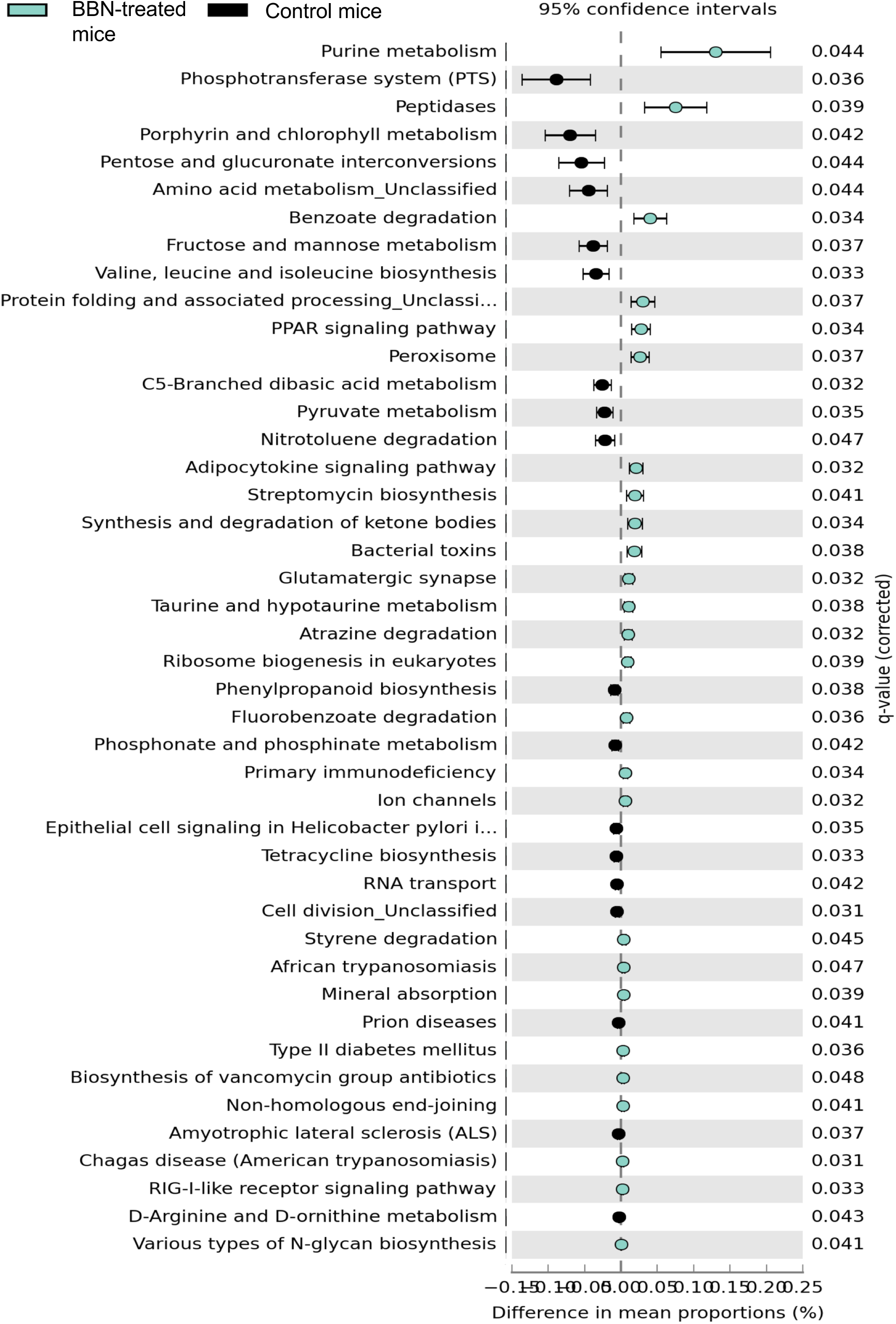
Inferred functional profiles of BBN-treated Mice and Control Mice at Month 4. Inferred functional profiles of BBN-treated mice versus control mice at month 4. Multiple pathways are inferred to be differentially upregulated in either group. Pathways with black point are increased in the control mice, and those with the blue point are increased in the BBN-treated mice.

When comparing control mice to the individual BBN sub-divisions, there were no differences in the relative abundance of microbial groups between the control and either BBN-precancer or BBN-cancer at any time point. However, BBN-normal mice had significantly higher proportions of *Gardnerella*, *Haemophilus*, *Bifidiobacterium*, and *Ureaplasma*, and significantly lower proportions of *Actinomyces* compared to the control mice at month 4. (Figure 5B) PCA plots demonstrated clustering of both the BBN-normal, BBN-precancer and the BBN-cancer group at month 4, which is distinct from their respective baseline urinary microbiomes. The control mice have urinary microbiomes at both month 0 and month 4 that cluster together, which is disparate from the month 4 cluster of the BBN mice. (Figure 3) There were no differences between controls and any individual BBN sub-division by PICRUSt analysis.

## Discussion

Here, we show that there are distinct urinary microbial profiles in mice exposed to 4 months of BBN compared to control mice. We report significant differences in several bacteria between the BBN-treated mice and control mice at month 4, but not in previous months. Further, we demonstrate stability in the urinary microbiome of control mice over a 4-month period.

While the overall shift in the composition of the microbial community is likely significant, there are also implications stemming from the specific bacterial operational taxonomic units (OTUs) that differ between these two groups. There were significantly higher proportions of *Gardnerella* and *Bifidobacterium* in the BBN-treated mice compared to control mice at month 4. *Gardnerella* has also been associated with cervical cancer. Women with cervical cancer had a higher incidence of *Gardnerella* in their vaginal flora compared with women with benign gynecological disease.[36] Further, there is a correlation between presence of *Gardnerella* and dysplastic changes in women undergoing screening for cervical cancer,[37] although more recent work in women with cervical cancer have not found this association.[38] In addition to *Gardnerella*, there were also significantly higher proportions of *Bifidobacterium* in the BBN-treated mice at month 4. While *Gardnerella* is associated with the development of cancer, *Bifidobacterium* has been shown to exert antitumor immunity in various mouse models.[39] Further, *Bifidobacterium* was also one of several components of the microbiota in patients with metastatic melanoma who responded to immunotherapy.[40] Similarly, patients with an increased risk of advanced colorectal cancers demonstrated a significantly lower proportion of *Bifidobacterium* in their fecal microbiota compared to patients with normal risk.[41] The significant increase in proportion of *Bifidobacterium* in the cohort of BBN-treated, but not control, mice suggests its involvement in oncogenensis. However, as we are underpowered to investigate the various sub-groups of mice (i.e. pre-cancer versus cancer versus normal), it is possible that the increase in *Bifidobacterium* in the BBN mice is seen only in the mice who were exposed to BBN, but did not develop cancer. Further work is needed to fully investigate the specific role of *Bifidobacterium* in bladder cancer.

We identified differentially increased proportions of bacteria in the BBN-treated mice that had histologically normal bladders, but with neither precancer nor cancer. One potential explanation is the low numbers of mice in the precancer and cancer sub-groups; Three mice had histology consistent with precancerous lesions, and only two mice developed cancer. These numbers may be too small to identify a difference in the proportion of individual members of their respective microbial communities. However, given the separation of clusters seen on the PCA chart, it is likely that we are underpowered to find a difference. Interestingly, one BBN-treated mouse with a distinct urinary microbiome at month 0 (Figure 1) went on to develop cancer. The predominant organisms in the baseline urinary microbiome of this mouse were largely different from those in the remaining mice. The most common organism in the urinary microbiome of the outlying mouse was *Rubellimicrobium*, followed by *Escherichia*, and then *Kaistobacter.*

*Escherichia* was highly prevalent in controls and BBN-treated mice, thus its role in this model is difficult to discern. In contrast, *Rubellimicrobium* and *Kaistobacter*were highly prevalent only in the outlier. *Rubellimicrobium* is a gram negative organism, found mostly in soil.[42] There is little in the literature about this organism, with no reports of association with diseases. Similarly, *Kaistobacter* is also found within the soil, with no reported association with specific disease states.[43] The implications of the presence of these organisms is unknown given the paucity of data within the literature. As this data is from a single mouse, we were not powered to examine the effects of alterations in the baseline microbiome with increased risk of bladder cancer. However, this hypothesis warrants further exploration in future work.

The proposed role of the urinary microbiome in bladder cancer raises the potential of the utility of therapeutic probiotics. However, this is not a novel idea. Indeed, the one of the most effective agents used for the management of bladder cancer is the intravesical administration of Bacillus Calmette-Guérin (BCG). Although not fully elucidated, the proposed mechanism by which BCG exerts an anti-tumor effect is thought to be due to activation of the immune system and immune-mediated cytotoxicity, as well as potential cytotoxic effects of BCG itself.[44] Other bacterial agents have also shown to have therapeutic potential in bladder cancer. *Lactobacillus casei,* Shirota strain, has demonstrated anti-tumor properties in several murine models of cancer,[45,46] with comparable effects to BCG. The hypothesized mechanism by which *L. casei* exerts its anti-tumor effect is stimulation of macrophages to produce cytokines with anti-tumor properties, such as IL-12 and tumor necrotic factor alpha.[47] The successful use of these microbiome-modulating agents in the treatment of bladder cancer provides further support for the existence of the bladder oncobiome.

While the role of asymptomatic bacteriuria in the urinary microbiome is unclear, data suggests that recurrent asymptomatic bacteriuria offers a protective effect against bladder cancer.[12] The mechanisms by which epithelial cells recognize pathogenic versus commensal organisms may offer a partial explanation for this observation, and allow for a more complete understanding of the role of the urinary microbiome in the pathogenesis of bladder cancer. The epithelial response to commensal organisms includes inhibition of the inflammatory response through the NF-κB pathway.[48] This anti-inflammatory effect on epithelial tissues has been demonstrated by a variety of commensal bacteria, including strains of *Lactobacillus*,[49] *Bifidobacterium*,[50] and *Fusobacterium*[51]. Conversely, pathogenic bacteria – specifically uropathogenic *E. coli* (UPEC) – can stimulate the inflammasome in urothelial cells, whereas non-pathogenic strains of *E. coli* do not.[52] Taken together, these data suggest that the response of host tissue to specific bacteria within the microbiome may modulate cancer-associated inflammation. Indeed, dysregulation of the extracellular matrix (ECM), which can occur as a result of age or a variety of diseases, plays a critical role in tumorigenesis through generation of a tumorigenic environment, including facilitation of angiogenesis and inflammation.[53] Further, inflammation induces ECM remodeling and generation of reactive oxygen species, leading to DNA damage, mutations, and ultimately oncogenesis.[54] There are, however, additional hypotheses as to how the urinary microbiome influences the development of bladder cancer. These include the metabolism of pro-carcinogenic environmental toxins, the direct effect of various bacterial virulence factors, and directly genotoxic bacterial metabolites (reviewed by Xu et al. [14] and Whiteside et al. [15]).

Another potential mechanism through which changes in the microbiome can lead to development of cancer is through the presence of biofilms. Biofilms have been implicated in the development of colon cancer: Dejea et al. demonstrated an association between colon cancer and the presence of dysbiotic biofilms. Their data also suggests that the presence of a biofilm in healthy controls is associated with procarcinogenic inflammation.[55] These authors hypothesize that biofilm formation increases permeability of the colonic epithelium that enables bacteria to directly interact with the unprotected epithelium surface, which facilitate development of procarcinogenic inflammation.[55] Given this hypothesized role of biofilms in colon cancer, and the potential association between recurrent UTIs and bladder cancer, it is plausible that biofilms also mediate procarcinogenic inflammation in the bladder. Indeed, species of *Gardnerella*, which were significantly increased in BBN-treated mice at month 4 compared to controls, can form biofilms,[56] and in this capacity may play a role in oncogenesis. Biofilms are also known to occur in the setting of UTI. *E. coli* forms intracellular biofilms,[57] while other bacteria form biofilms associated with indwelling devices, such as urinary catheters.[58] Our data provide some support for this: bacterial gene pathways predicted to be significantly expressed at increased levels in the urinary microbiota of the BBN mice at 4 months include those that would increase inflammation of the epithelium, e.g., peptidase and toxin production.

There are several other examples of cancers associated with changes in their respective microbiomes. Patients with grade 4 oral squamous cell carcinoma have significant differences in their oral microbiome compared to healthy controls, with changes in community complexity, as well as composition. Further, the proportion of *Fusobacterium* increased while the proportions of *Streptococcus*, *Haemophilus*, *Porphyromonas*, and *Actinomyces* progressively decreased with increased disease severity, with the largest differences seen in stage 4 carcinoma.[59] Similar work has been conducted in cervical cancer, where women with increasingly severe grades of carcinoma *in situ* have decreasing proportions of *Lactobacillus* in their vaginal microbiomes.[60] Patients with cirrhosis who progress to develop hepatocellular cancer have different gut microbiomes than those who do not develop cancer.[61] These data suggest that altered microbial profiles exist in a variety of oncologic conditions, including bladder cancer.

Limitations of this study include the small number of mice that developed bladder cancer. Further, the exposure to BBN may have affected our results independent of the changes in histology. Finally, the use of 16S rRNA sequencing, rather than whole genome sequencing, limits our ability to make inferences about the metagenomes in bladder cancer. Most of the OTUs we identified in this work were classified to the genus level, and some only to the family level. Resolving the data to the species or even strain level would be more informative. Future work will include a larger number of mice, the use of additional mouse models of bladder cancer to ensure that these results are due to the presence of bladder cancer rather than result of exposure to BBN, and use whole genome sequencing.

## Conclusion

Mice exposed to 4 months of BBN, a bladder-specific carcinogen, have distinct urine microbial profiles compared to control mice. This suggests that there are specific urine microbial profiles associated with the development of bladder cancer, which we propose represents the existence of a “urinary oncobiome”.

